# Comparison of genomic DNA isolation methods in *Pereskia aculeata Mill*. (Ora-pro-nóbis)

**DOI:** 10.1101/2023.10.09.561527

**Authors:** Joselina Barbosa da Silva Neta, Amaro Antônio Silva Neto, Thaís Correia Magalhães, Michely Correia Diniz

**Affiliations:** Universidade Federal do Vale do São Francisco – UNIVASF, Agricultural Sciences Campus -Rodovia BR 407, 12 Lot 543, Irrigation Project -Nilo Coelho - S/N C1-CEP: 56300-990 Petrolina/PE/Brazil

**Keywords:** UFP, DNA protocols, Plants

## Abstract

In Brazil, it is estimated that at least 10% of the native flora, which corresponds to about 4 or 5 thousand plant species, are used for food purposes. Among these species, *Pereskia aculeata* Mill, known as ora-pro-nóbis and belonging to the Cactaceae family, stands out. It is considered a Unconventional Food Plant (UFP) and a vegetable of traditional use, playing an important role in the diet and economy of rural and urban families. Due to their high nutrient content, they have both economic and social significance, being a target of interest for scientific research. Despite the potential for human use of this species, there is a lack of knowledge about its diversity and genomic structure, resulting in few genetic and molecular studies that can contribute to the conservation and improvement of this species in general. It is essential to acquire information about its genomic structure, as well as to carry out studies related to the comparison, selection and optimization of protocols for obtaining quality DNA and in sufficient quantity, in order to guarantee the success of molecular and evolutionary analyses. The aim of this research was to evaluate qualitatively and quantitatively methods of isolation of genomic DNA from *Pereskia aculeata* Mill. Significant variations were observed in the DNA extracted and in the purity observed between the different protocols tested, with the isolation by the protocol described by Sambrook et al. (1989) being the most efficient in terms of concentration and purity, followed by the modified method proposed in this work. This work is a pioneer in comparing methods for isolating DNA from *Pereskia aculeata* Mill. in terms of quality and quantity. In addition, it was found that there is still a scarcity of molecular studies on this species, both worldwide and in Brazil, which indicates the existence of a promising gap for the development and realization of scientific research in this field.

## 1. INTRODUCTION

It is estimated that around 400,000 plant species are known in the world and of these, 7,039 are used for food purposes^1^. Many of these are defined as “Unconventional Food Plants” (UFPs) with economic and ecological potential, and their roots, stems, leaves, flowers or fruit can be used for human consumption^2,3^.They are plants typical of some regions that are not included in the daily menu, including native, exotic, cultivated and spontaneous species^2^. In Brazil, at least 10% of the native flora (around 4 or 5 thousand plant species) is used for food purposes^4^.

Among them is *Pereskia aculeata* Mill (Ora-pro-nóbis) (OPN), belonging to the Cactaceae family and widely distributed, the species is native to the American tropics, occurring in 12 Brazilian states and in various types of environments^5,6^. It is a perennial plant with succulent and lanceolate leaves, its fruits are small yellow berries are small yellow berries, its flowers are small and white and on its stem there are (false thorns)^7^. This species is defined as UFPs and a vegetable of traditional use, contributing to the diet and family economy of rural and urban populations^8^.

The plants of this species are of great economic and social importance due to their high nutrient content, with notable levels of protein, fiber, iron, calcium, vitamins (A and C) and bioactive compounds such as carotenoids, as well as the absence of toxicity and large amount of mucilage in its leaves and stems, arousing interest in the food and pharmaceutical industries, in addition to its use in traditional medicine^9^. Due to its biological and nutritional characteristics, OPN is of great relevance to scientific research^10^.

Although this species has great potential for human use, little is known about its diversity and structure at genome level, and consequently there are few genetic and molecular studies that can contribute to conservation and improvement actions for this species. Until the mid-1960s, the evaluation of genetic diversity was based on morphological characters, generally phenotypes that were easy to identify visually. However, with the development of new technologies in the field of molecular biology, especially the polymerase chain reaction (PCR), various molecular methods have been created to detect existing variability directly at the DNA level^11,12^.

The isolation of DNA from plant tissues plant tissues is one of the most important stages and requires specific protocols for obtaining pure, intact DNA that enables the genomic material to be amplified. However, OPN is a species rich in secondary metabolites, polysaccharides and mucilage, which make it difficult to isolate DNA in quantity and quality for molecular analysis^13-15^.

For these reasons, knowledge of the genomic structure of *Pereskia aculeata* Mill. (Ora-pro-nóbis), as well as conducting studies associated with the comparison, selection and optimization of protocols for obtaining DNA of sufficient quality and quantity, in order to guarantee the success of studies for the main molecular and evolutionary analyses.

The aim was to evaluate qualitatively and quantitatively methods of isolation of genomic DNA from *Pereskia aculeata* Mill.

## 2. METHODS

### Leaf sample

The samples of *Pereskia aculeata* Mill. were collected on the agricultural sciences campus of the Federal University of the São Francisco Valley (CCA/UNIVASF) (−9.325195, -40.549271) and in a popular market, located in the municipality of Petrolina in the state of Pernambuco, Brazil. The access to genetic heritage was registered in Sisgen under number A5DA19B.

For the plant material, young, fresh, healthy and dirt-free leaves were collected. They were placed in plastic bags, packed in Styrofoam with ice and transferred to the Genetics and Biotechnology (CCA/UNIVASF) and Plant Biotechnology (EMBRAPA) laboratories, where they were stored in a -20°C freezer and -80°C ultrafreezer, respectively, until DNA isolation.

### Isolation of genomic DNA

The genomic DNA isolation tests were carried out according to four previously published methods (See Appendices 1,2,3 and 4) based on a single protocol^16^ (P1) with modifications described in Table 1.

**Table 1.** Variations in DNA isolation methods.

| Protocols | Main steps involved in the procedure |
| --- | --- |
| P1 | CTAB 2%*+ chloroform/isoamyl alcohol 24:1+ Isopropanol+ Ethanol 70%+ Water ultra pure; |
| P2 | Pre-wash with sorbitol buffer*+ CTAB 3%*+ NaCl 3M+ chloroform/alcohol isoamyl 24:1+ Isopropanol/sodium acetate+ 70% ethanol+ ultrapure water; |
| P3 | CTAB 2%*+ Phenol/chloroform/isoamyl alcohol 25:24:1+ Isopropanol+ Ethanol 70%+ Ultra pure water; |
| P4 | Pre-wash with sorbitol buffer*+ CTAB 3%*+ NaCl 3M+ Phenol/chloroform/isoamyl alcohol 25:24:1+ Isopropanol/sodium acetate+ Ethanol 70%+ Ultra pure water. |

\* Addition of β-mercaptoethanol at the time of use. Source: Author (2023).

In the second protocol^17^ (P2), the main modification proposed was the addition of a pre-wash with sorbitol buffer applying 3% CTAB with a high salt content, to reduce the number of polysaccharides in the extracted DNA. The third protocol^18^ (P3) was modified by adding phenol/chloroform/isoamyl alcohol 25:24:1 in the purification stage. Finally, the fourth protocol (P4) was modified by combining protocols P2 and P3, modified by pre-washing with sorbitol buffer and the addition of phenol-chloroform. Six replicates were carried out for each protocol.

### Analysis of total isolated DNA

After the purification stage, the nucleic acids were assessed for their integrity by electrophoresis on a 1% (w/v) agarose gel for better attribution of the results, stained with *GelRed® (Biotium)* and run in a 0.5X TBE solution (Tris-Acid-Boric-EDTA) for 60 minutes at 80 V. The results were compared with the molecular weight marker (Lambda DNA; *Invitrogen*) at concentrations of 25, 50 and 100 ng/μL. The gel was viewed under UV light and recorded using the *LPix Image* photodocumentation system version 2.7 (Loccus Biotecnologia®).

Purity determination and concentration estimation was carried out using a NanoDrop® spectrophotometer (*Thermo Fisher Scientific*), measuring the absorbance in contrast to an ultra-pure water sample in the ultra-violet range at 260 (DNA quantification) and 280 (protein quantification). The degree of purity was established using the A260/A280 ratio.

### Comparison of time for the different DNA isolation protocols

The time taken to isolate DNA using each method was determined considering the average time taken to carry out the main steps: isolation, which involves incubation in a water bath with the extraction buffer, precipitation, cleaning with CIA or FCIA until the pellet is obtained, disregarding previous steps for preparing the plant material and subsequent steps for cleaning. Quick actions such as adding and mixing reagents and using the vortex were not considered.

## 3. RESULTS

### Comparison of the efficiency of DNA isolation techniques

Preliminary analysis of the “quality” aspect was carried out using electrophoresis in a 1% agarose gel (Figure 1). After electrophoretic migration, the extracted DNA appeared on the agarose gel in the form of well-delineated bands with medium fluorescence, a result of the good performance of the isolation protocols for most of the samples. The image also showed a good amount of DNA excreted by the bands at concentrations ranging from 25 to 100 ng/μL (compared to the DNA λ standard), and satisfactory quality for the P4 protocol, demonstrated by the sharpness and lack of residues in the migration channel.

**Figure 1.**
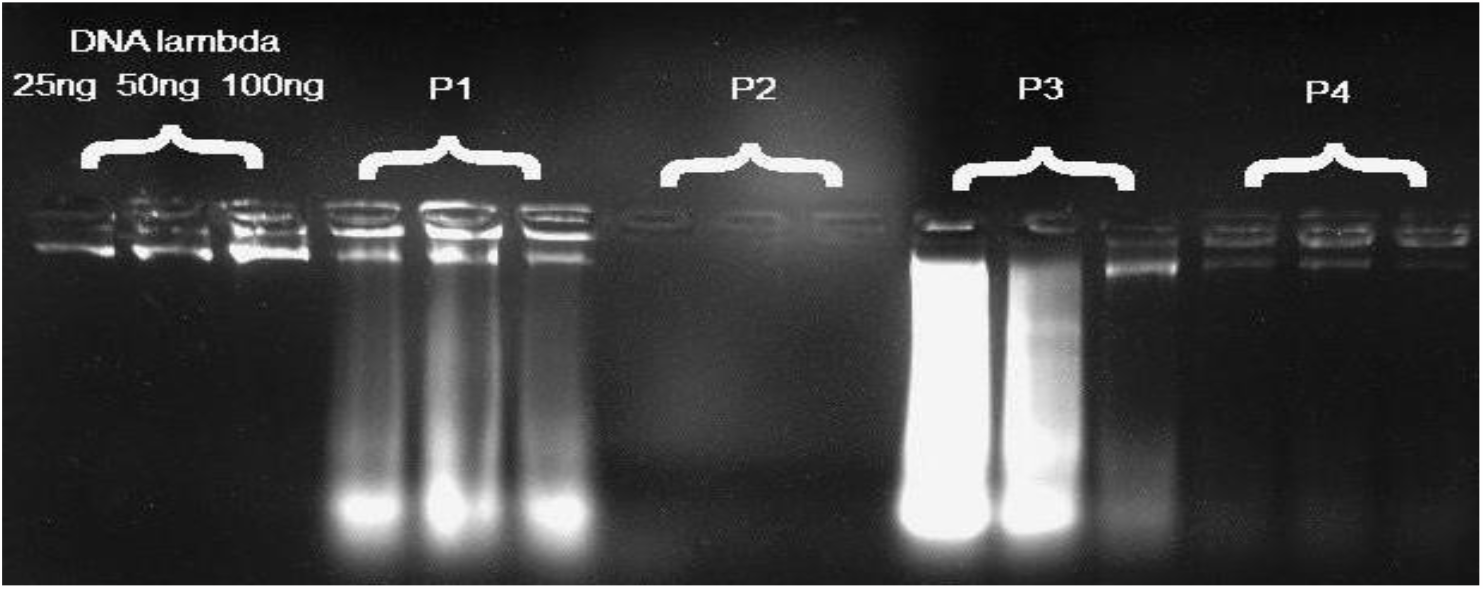
Genomic DNA from *Pereskia aculeata* Mill. DNA λ at different concentrations (ng/μ). P1: method proposed by Doyle & Doyle (1987). P2: protocol modified by Inglis *et al*. (2016). P3: protocol based on Sambrook *et al*. (1989). P4: protocol modified by combining protocols P2 and P3. Source: Author (2023).

When analyzed separately, the images show the P4 protocol with the best results, followed by P1. Protocol P2 showed no visible bands with any sample. Although protocol P2 showed no visible bands on the gel after electrophoresis, and protocols P1 and P3 showed signs of DNA degradation, spectrophotometry showed that genetic material was present in these samples based on the concentrationvalues obtained.

The readings and concentration of DNA in the samples, as well as the A260/A280 ratio, are shown in Table 2, where the spectrophotometric data shows genetic material in all the samples, but in varying concentrations.

**Table 2.** Average values for estimating DNA purity and concentration by spectrophotometric reading(A260/280 ratio).

| Protocol | A260/A280 | Concentration (ng/μL) |
| --- | --- | --- |
| P1 | 1,59 | 400 |
| P2 | 1,33 | 712,58 |
| P3 | 2,03 | 1.897,91 |
| P4 | 1,40 | 827 |

Source: author (2023).

The protocols P1, P2 and P4 presented samples that are not considered “pure”, as well as showing a yellow or brown color pattern, probably indicating contamination with oxidized polyphenols (Figure 2).

**Figure 2.**
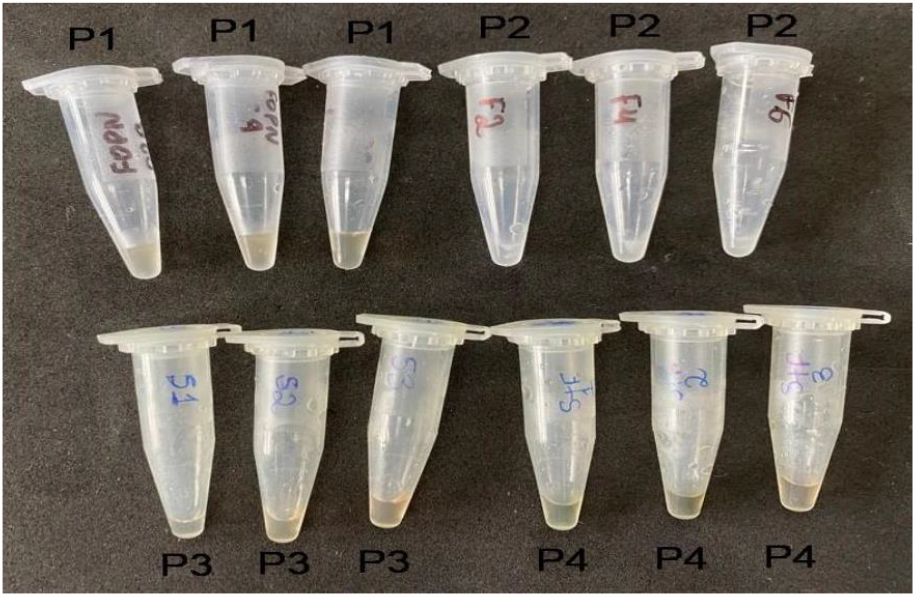
Resuspended DNA samples isolated from *Pereskia aculeata* P1: method proposed by Doyle & Doyle (1987). P2: protocol modified by Inglis *et al*. (2016). P3: protocol based on Sambrook *et al*. (1989). P4: protocol modified by combining protocols P2 and P3. Source: Author (2023).

The results of the absorbance ratio of 260/280 showed that the genetic material obtained in protocol P3 met acceptable standards of contamination by secondary compounds and had preserved integrity.

By comparison, it was found that the concentration value of all the DNA samples measured by spectrophotometer was much higher than by agarose gel electrophoresis. The spectrometric estimates related to the concentration of genomic DNA obtained ranged from 400 ng/μL for P1 to 1,897.91 ng/μL for P3. This range in DNA concentration estimates demonstrates the influence that the choice of method adopted for DNA isolation has on *P. aculeata* Mill.

### Comparison of DNA isolation times

The DNA isolation time of the protocols tested ranged from 140 to 330 minutes, with protocol P2 being the fastest and P3 the longest (Table 3).

**Table 3.** Time spent, in minutes, on the main stages of the DNA isolation protocols studied. P1-Isolation: incubation in a water bath; Purification: three cleanings with CIA (homogenization and centrifugation); Precipitation: centrifugation to precipitate the pellet. P2-Isolation: washing in sorbitol and incubation in a water bath; Purification: centrifugation to separate the supernatant from the DNA; Precipitation: centrifugation to precipitate the pellet. P3 Isolation: incubation in a water bath; Purification: cleaning with FCIA three times (homogenization and centrifugation); Precipitation: centrifugation to precipitate the pellet. P4-Isolation: washing in sorbitol and incubation in a water bath; Purification: centrifugation to separate the supernatant from the DNA; Precipitation: centrifugation to precipitate the pellet.

| Procedure | Protocols |  |  |  |
| --- | --- | --- | --- | --- |
|  | P1 (min) | P2 (min) | P3 (min) | P4(min) |
| Isolation | 60 | 70 | 60 | 70 |
| Purification | 30 | 10 | 30 | 30 |
| Precipitation | 120 | 60 | 240 | 60 |
| Total time | 210 | 140 | 330 | 160 |

All the protocols tested use 30 to 60 minutes of incubation in a water bath. For protocols P2 and P4, there is a 10-minute increase in the isolation stage due to pre-washing with sorbitol buffer. In addition, protocols P1 and P3 require more than one step of cleaning with CIA, which considerably increases their execution time, except for protocols P2 and P4, which do not need to repeat this step.

Despite showing better results in terms of DNA isolation quantity and quality, the total isolation time of protocol P3 was even longer than the others due to the precipitation step, since in the former the samples are kept at - 20 ºC for four hours to precipitate the DNA, which is not necessary in the other protocols tested.

Although isolation using protocol P2 had a higher cost in the cell lysis stage, its total cost was lower when compared to the others since there were no repetitions between the stages. Furthermore, even when using sorbitol and CTAB 3% (P2 and P4), the cost was higher for the protocols that only used CTAB 2% (P1 and P3).

Purification accounted for the highest total costs in protocols P1, P2 and P3, since in protocols P1 and P3 three cleansings were carried out with CIA and FCIA, respectively, which increases the cost of these reagents. Precipitation was the least costly step, except for protocol P4, which accounted for 51.83% of the total costs, which can be explained by repeating the ethanol washing step once more. Therefore, protocol P1 proved to be the costliest and time consuming, while the P2 protocol proved to be the fastest and least costly when compared to the other protocols.

## 4. DISCUSSION

The A260/280 absorbance estimate is used to assess the purity of nucleic acids (DNA and RNA), making it possible to infer the presence of protein and phenol, among other possible contaminants. The minimum acceptable value found in the literature for the absorbance equation (260nm/280nm) is between 1.8 and 2 and indicates pure DNA, that is, lower values indicate protein contamination while higher values indicate phenol contamination^19^.

A challenge for subsequent molecular manipulations is often observed in samples that have a high content of oxidized polyphenols, which are known to bind irreversibly to DNA^20^. In this sense, the addition of β-mercaptoethanol serves to inhibit the oxidation of polyphenols. Polysaccharide contamination in the samples manifested itself as a gelatinous, sticky substance after isopropanol precipitation and centrifugation, making resuspension and pipetting difficult.

The DNA was purified using buffered phenol, which efficiently denatures the proteins. Chloroform also plays a role as a denaturing agent for the proteins present in the sample. The mixture of phenol and chloroform is highly effective in deproteinization, based on the hydrophobic property of proteins, which have an affinity for organic solvents^21^.

CTAB in saturated NaCl was initially used to preserve leaves^22^. Since then, some modifications to the method have been tested and published. Some authors suggested homogenizing the samples in sorbitol wash buffer before isolation with CTAB^23,24^ Some studies obtained good yields of high purity DNA by adding a pre-wash step with sorbitol combined with a higher concentration of CTAB with a high salt content, tested on several different plant species, including *P. aculeata*. This pre-wash aims to remove interfering metabolites such as polyphenols and polysaccharides from macerated tissues^17^.

One study carried out a comparison between two DNA isolation protocols using CTAB at concentrations of 2% and 5%. It was observed that the buffer containing CTAB 5% was efficient in isolating DNA from Anacardium giganteum (Cashew) leaves^25^. Similarly, another study found that the buffer containing CTAB 2% was not efficient, while the buffer containing CTAB 5% was efficient in isolating DNA from the *Curcuma longa* species. Unlike the results obtained in the present study, the different concentrations of CTAB did not prove to be a determining factor in resulting in a high concentration of genomic DNA (ng/μl), as can be seen in the electroforese image (Figure 1)^26^.

DNA samples with a low concentration may not hinder the reproducibility of PCR reactions, but they can make it difficult to properly balance the reaction mixture^15, 27^. When samples with a low DNA concentration are obtained, it is recommended that a new isolation attempt is made to optimize time and reduce costs related to PCR replications.

Adjusting the concentration of DNA is extremely important, since too much can result in complete failure of the reaction due to the high concentration of aggregated impurities and can also lead to electrophoretic profiles with drags and poorly defined bands^28^. On the other hand, a low concentration of DNA can result in irregular amplifications or even a lack of amplification of certain fragments, with electrophoresis profiles that are not reproducible ^29,30^.

## 5. CONCLUSION

This work is a pioneer in comparing methods for isolating DNA from *Pereskia aculeata* Mill. in terms of quality, quantity, and cost, as well as describing information about its genome. The isolation of genomic DNA was influenced by the type of reagent used in the purification stage, with phenol:chloroform:isoamyl alcohol 25:24:1 having a positive correlation, so the results of this study indicate that the most suitable protocol for extracting genomic DNA from *P. aculeata* Mill. is the P3 protocol, as it presents more satisfactory data.

These results open a promising prospect for the use of the DNA samples collected in the application of molecular techniques such as amplification, cloning and sequencing. In this context, the use of DNA becomes viable for more advanced studies, despite the difficulties encountered during the isolation of ora-pro-nóbis genetic material.

## APPENDICE 1 PROTOCOL FOR ISOLATING DNA FROM PLANT TISSUE: P1-(DOYLE & DOYLE, 1987)

Procedure:

a) Weigh out 300 mg of plant tissue (the amount of plant tissue can be up to 3 g).
b) Macerate with liquid nitrogen until the sample is pulverized.
c) Place the sample in a 2 mL microtube and add 700 μL of plant extraction buffer solution preheated to 65° C containing 2-mercaptoethanol and homogenize for ten minutes by inversion.
d) Incubate the samples in a water bath at 65ºC for one hour, homogenizing every 15 minutes.
e) Remove from the water bath and allow the temperature of the sample to equalize to room temperature.
f) Add 700 μL of chloroform and isoamyl alcohol solution (24:1) and homogenize gently.
g) Centrifuge at 13000 rpm for ten minutes at room temperature to separate the organic and aqueous phases.
h) Remove the aqueous phase (top) into a new microtube, avoiding touching the inter-surface with the tip.
i) Repeat the extraction with a solution of chloroform and isoamyl alcohol (24:1) one or two more times, considering that a greater number of extractions can purify the DNA more, but with some loss.
j) After the final removal of the supernatant, add isopropanol (ice-cold), equivalent to approximately 2/3 of the volume collected. Homogenize gently.
k) Keep at -20ºC for at least two hours and centrifuge at 13000 for ten minutes.
m) Discard the supernatant.
n) Wash the pellet once with 70% ethanol.
o) Wash the pellet again with 95% ethanol.
p) Leave to dry for up to an hour.
q) Resuspend in 50 to 100 μL of autoclaved milli Q water or TE buffer (RNAse can be added at a final concentration of 10 μg/mL of sample).
r) Keep at -20ºC.

## APPENDICE 2 PROTOCOL FOR ISOLATING DNA FROM PLANT TISSUE: P2-(INGLIS *et al*., 2016)

Procedure:

a) Add an excess of sorbitol wash buffer to fill the sample tubes containing macerated plant material to approximately ¾ of the capacity (0.9 to 1.5 ml, depending on the tubes used). Cap the tubes and shake them in the bead mill, either manually or using the vortex. Inspect to confirm suspension of the powdered material and shake again if necessary.
b) Centrifuge at 2,500 x g for five minutes at room temperature.
c) The supernatant is decanted or aspirated from the samples and discarded.
d) (Optional) Repeat the sorbitol wash if the supernatant from step 3 is too turbid or viscous. Resuspend and centrifuge as before. Discard the supernatant. In general, one sorbitol wash cycle is sufficient, but it can be repeated, especially with samples that are particularly rich in polysaccharides or tannins, judging by the clarity and color of the supernatant after centrifugation.
e) Add the 3% CTAB extraction buffer preheated to 65° C to the sample tubes (approximately ½ of the sample tube capacity or 500 to 700 μl, for example, when using 1.1 ml microtubes in 96 wells).
f) Resuspend the macerated samples by shaking them for five seconds in the ball mill or by stirring them in a vortex. The ball bearings that remain in the tubes greatly aid the mixing process of the macerate.
g) Incubate the samples at 65 °C for a minimum of 30 minutes and up to 60 minutes, mixing them by inversion every ten minutes. Remove from the water bath/oven and allow the samples to cool at room temperature for five minutes.
h) Add CIA 24:1 to the sample tubes (700 μl or fill to approximately 4/5 of the tube capacity). Shake the tubes vigorously for 10 seconds. This can be done efficiently using the ball mill, if desired.
i) Centrifuge the samples at 2,500 x g for ten minutes at room temperature.
j) The nucleic acids are precipitated by adding 1/10 volume of 3 M sodium acetate pH 5.2 and 0.66 volume of cold isopropanol (stored at -20 °C). Mix by inverting ten times and keep at -20 °C for at least one hour.
k) Centrifuge at 13,000 x g for 10 minutes at room temperature
l) Carefully decant the supernatants and drain the tubes, placing them inverted on paper towels.
m) Wash the pellets by adding 1 ml of 70% ethanol. Centrifuge at 13,000 x g for 10 minutes. Carefully remove the supernatants by aspiration to avoid losing the nucleic acid pellet.
n) Dry the opened tubes at room temperature for approximately one hour or vacuumdry for 10 minutes.
o) Gently suspend the pellets in 100 μl of TE containing 0.1 mg ml-1 RNAse without DNase and incubate the tubes at 37 °C for 30 minutes.

## APPENDICE 3 PROTOCOL FOR ISOLATING DNA FROM PLANT TISSUE: P3 (SAMBROOK *et al*., 1989)

Procedure:

a) Weigh out 300 mg of plant tissue (the amount of plant tissue can be up to 3 g).
b) Macerate with liquid nitrogen until the sample is pulverized.
c) Place the sample in a 2 mL microtube and add 700 μL of 2% CTAB plant extraction buffer solution and homogenize for ten minutes by inversion.
d) Incubate the samples in a water bath at 65ºC for one hour, homogenizing every 15 minutes.
e) Remove from the water bath and allow the temperature of the sample to equalize to room temperature.
f) Add 700 μL of phenol:chloroform:isoamyl alcohol solution (24:1) and homogenize gently.
g) Centrifuge at 9000g for ten minutes at room temperature to separate the organic and aqueous phases.
h) Remove the aqueous phase (top) into a new microtube, avoiding touching the inter-surface with the tip.
i) Repeat steps f to h 1 or 2 times.
j) After the final removal of the supernatant, add isopropanol (ice-cold), equivalent to approximately 2/3 of the volume collected. Homogenize gently.
k) Keep at -20ºC for at least four hours and centrifuge at 13000 for ten minutes.
m) Discard the supernatant.
n) Wash the pellet once with 70% ethanol and wash the pellet again with absolute ethanol.
p) Leave to dry for up to an hour.
q) Resuspend in 50 to 100 μL of autoclaved milli Q water or TE buffer (RNAse can be added at a final concentration of 10 μg/mL of sample).
r) Incubate for 4 hours at 37°C or store at -20°C.

## APPENDICE 4 PROTOCOL FOR ISOLATING DNA FROM PLANT TISSUE: P4 (MODIFIED BY THE AUTHOR, 2023)

Procedure:

a) Add an excess of sorbitol wash buffer to fill the sample tubes containing macerated plant material to approximately ¾ of the capacity (0.9 to 1.5 ml, depending on the tubes used). Cap the tubes and shake them in the bead mill, either manually or using the vortex. Inspect to confirm suspension of the powdered material and shake again if necessary.
b) Centrifuge at 2,500 x g for five minutes at room temperature. The supernatant is decanted or aspirated from the samples and discarded. (Optional) Repeat the sorbitol wash if the supernatant from step 3 is too turbid or viscous. Resuspend and centrifuge as before. Discard the supernatant. In general, one wash cycle with sorbitol is sufficient, but it can be repeated, especially with samples that are particularly rich in polysaccharides or tannins, judging by the clarity and color of the supernatant after centrifugation.
c) Add the 2% CTAB extraction buffer preheated to 65° C to the sample tubes (approximately ½ of the sample tube capacity or 500 to 700 μl, for example, when using 1.1 ml microtubes in 96 wells).
d) Resuspend the macerated samples by shaking them for five seconds in the ball mill or by stirring them in a vortex. The ball bearings that remain in the tubes greatly aid the mixing process of the macerate.
e) Incubate the samples at 65 °C for a minimum of 30 minutes and up to 60 minutes, mixing them by inversion every ten minutes. Remove from the water bath/oven and allow the samples to cool at room temperature for five minutes.
f) Add FCIA 25:24:1 to the sample tubes (700 μl or fill to approximately 4/5 of the tube capacity). Shake the tubes vigorously for 10 seconds. This can be done efficiently using the ball mill, if desired.
g) Centrifuge the samples at 2,500 x g for ten minutes at room temperature.
h) Carefully remove the upper aqueous phase with a pipette. Transfer to a new tube, carefully avoiding disturbing the debris between the phases.
i) The nucleic acids are precipitated with an equal volume of cold isopropanol (stored at -20 °C). Mix by inverting ten times and keep at -20 °C for at least one hour.
j) Centrifuge at 13,000 x g for 10 minutes at room temperature. Carefully decant the supernatants and drain the tubes, placing them inverted on paper towels.
k) Wash the pellets by adding 1 ml of 70% ethanol. Centrifuge at 13,000 x g for 10 minutes. Carefully remove the supernatants by aspiration to avoid losing the nucleic acid pellet.
l) Dry the opened tubes at room temperature for approximately one hour or vacuum dry for 10 minutes.
m) Gently suspend the pellets in 100 μl of TE containing 0.1 mg ml-1 RNase A without DNase and incubate the tubes at 37 °C for 30 minutes.

